# Rehydration of freeze substituted brain tissue for pre-embedding immunoelectron microscopy

**DOI:** 10.1101/2021.12.14.472089

**Authors:** Janeth Pérez-Garza, Emily Parrish-Mulliken, Zachary Deane, Linnaea E. Ostroff

## Abstract

Electron microscopy (EM) volume reconstruction is a powerful tool for investigating the fundamental structure of brain circuits, but the full potential of this technique is limited by the difficulty of integrating molecular information. High quality ultrastructural preservation is necessary for EM reconstruction, and intact, highly contrasted cell membranes are essential for following small neuronal processes through serial sections. Unfortunately, the antibody labeling methods used to identify most endogenous molecules result in compromised morphology, especially of membranes. Cryofixation can produce superior morphological preservation and has the additional advantage of allowing indefinite storage of valuable samples. We have developed a method based on cryofixation that allows sensitive immunolabeling of endogenous molecules, preserves excellent ultrastructure, and is compatible with high-contrast staining for serial EM reconstruction.

## Introduction

Electron microscopy (EM) is the only imaging technique that reveals complete subcellular structure, and is therefore indispensable for visualizing synaptic connectivity in the brain. Serial section EM volume reconstruction is a powerful tool for studying the effects of experience on synapse organization (Sorra and Harris, 1998; Popov et al., 2004; Ostroff et al., 2010) and high-throughput implementations have been used to map complete circuits and even entire insect brains (Anderson et al., 2011; Bock et al., 2011; Helmstaedter et al., 2013; Kasthuri et al., 2015; Zheng et al., 2018; Phelps et al., 2021). Methods for incorporating molecular information into EM reconstructions are very limited, which is a serious drawback given the importance of accounting for molecular heterogeneity among neurons (Yuste et al., 2020). The major barrier to combining immunolabeling with EM reconstruction is the low tolerance of the latter for compromised morphology or staining. Successful reconstruction requires the ability to readily and unambiguously follow small neuronal processes across hundreds or thousands of sections, and this in turn requires uniformly intact, strongly contrasted membranes and well-preserved, easily recognizable subcellular structures. Molecular labeling must therefore be accomplished without impairing morphology or interfering with the strong staining needed for volume reconstruction.

Brain tissue is typically prepared for serial EM by a combination of strong fixation and multiple heavy metal staining steps to maximize structural preservation and contrast (Sorra and Harris, 1998; Ostroff et al., 2010; Tapia et al., 2012; Hua et al., 2015; Genoud et al., 2018). These protocols invariably include osmium tetroxide, which provides essential membrane contrast but also prevents labeling of most protein antigens (Berryman and Rodewald, 1990; Phend et al., 1995). Large-scale EM reconstructions have incorporated on-section immunolabeling for amino acid and peptide neurotransmitters (Anderson et al., 2011; Shahidi et al., 2015), both of which are osmium-resistant. Most other targets require pre-embedding labeling, in which antibodies are applied before osmium staining (Sesack et al., 2006; Polishchuk and Polishchuk, 2019). A disadvantage to pre-embedding labeling is that reagents must access target molecules in fixed samples without the aid of ultrathin sectioning to expose the sample interior. Approaches to enhance antibody penetration include reducing or replacing the primary glutaraldehyde fixative (Somogyi and Takagi, 1982; King et al., 1983; Fulton and Briggman, 2021), permeabilizing the tissue with detergents or freeze-thaw cycles (Eldred et al., 1983; Pickel et al., 1986), and using extensive antibody incubation times (Fulton and Briggman, 2021), all of which compromise ultrastructure. Genetically-encoded tags that withstand osmium (Viswanathan et al., 2015) or minimize the need for reagent penetration (Schikorski et al., 2007; Martell et al., 2012; Cruz-Lopez et al., 2018) have been developed, but methods for detecting endogenous markers without ultrastructural damage are still needed.

Cryofixation by high-pressure freezing (HPF) produces superior ultrastructure to standard methods (McDonald and Auer, 2006; Vanhecke et al., 2008; Korogod et al., 2015), even in brain tissue that has been previously fixed with aldehydes (Sosinsky et al., 2008). HPF results in amorphous ice instead of the damaging hexagonal ice that forms during slower freezing (Moor et al., 1980) so that samples can be frozen without impairing ultrastructure with cryoprotectants (Gilkey and Staehelin, 1986). Subsequent dehydration at low temperature, called freeze substitution, preserves structure by minimizing extraction of sample molecules by the organic solvent (Weibull and Christiansson, 1986; Kellenberger, 1991). Unlike standard EM preparation protocols, which must be performed immediately after sample collection to prevent degradation of morphology, HPF allows samples to be stored indefinitely in liquid nitrogen. This is especially useful for valuable samples with unpredictable availability, such as human tissue, or for experiments where large numbers of samples must be collected at once, such as behavioral studies.

Combining HPF with pre-embedding immunolabeling and EM reconstruction would allow valuable samples to be stored until protocols are optimized and convenient, but only if it can be done without compromising ultrastructure. Both immunolabeling and multi-step heavy metal staining are done in aqueous environments, so high-pressure frozen tissue must be rehydrated after freeze substitution. The challenge with rehydration is that it cannot begin until the sample is warm enough to prevent ice crystal formation, but warm solvents can extract and distort membranes. Osmium tetroxide is typically included in freeze substitution media for this reason, even for samples fixed in aldehydes before HPF (Sosinsky et al., 2008). Rehydration after freeze substitution in low concentrations of osmium has been used for immunolabeling on cryosections (van Donselaar et al., 2007; Ripper et al., 2008) and cultured cells (Hess et al., 2018) and for enzyme cytochemistry on cultured cells (Robinson and Karnovsky, 1991). By substituting and warming samples in a mixture of uranyl acetate and glutaraldehyde, osmium-free rehydration with good ultrastructure has been achieved for immunolabeling of cultured cells (Twamley et al., 2021) and enzyme cytochemistry in nervous tissue (Tsang et al., 2018). Antibody penetration into brain tissue can be adversely affected by strong fixation, so we developed a protocol that combines high-pressure freezing with rehydration using minimal additional fixation to allow subsequent pre-embedding immunolabeling of endogenous molecules. The ultrastructural quality of samples prepared with our protocol is as good as or better than that achieved with standard protocols, even after immunolabeling.

## Results

### Ultrastructural preservation during rehydration with minimal fixation

Our first goal was to determine whether we could return high-pressure frozen brain tissue to a thawed, hydrated state without compromising ultrastructure or using fixatives that would preclude later immunolabeling. Excellent ultrastructure can be obtained by high-pressure freezing live brain tissue and performing chemical fixation during the freeze substitution step (Rostaing et al., 2006; Frotscher et al., 2014; Korogod et al., 2015; Tamada et al., 2020), which allows the artifacts of chemical fixation to be minimized. HPF samples must be no more than 100µm thick, however, and sectioning live brain tissue introduces substantial artifacts from ischemia and mechanical trauma (Fiala et al., 2003). High-pressure freezing after chemical fixation also preserves brain ultrastructure (Sosinsky et al., 2008) and allows for dissection under reproducible fixation conditions, so we chose this approach. Rats were perfused with a mixture of glutaraldehyde and paraformaldehyde, which has long been considered optimal for preservation of brain ultrastructure (Karnovsky, 1965; Schultz and Karlsson, 1965), followed by HPF of vibratome sections containing the amygdala.

In the only study reporting rehydration of brain tissue without osmium fixation, samples were freeze substituted and then warmed in a mixture of uranyl acetate and glutaraldehyde (Tsang et al., 2018). Uranyl acetate was removed at -30°, but glutaraldehyde was present throughout the full protocol. Seeking to minimize the time, temperature, and strength of additional fixation, we tested protocols in which any fixative used in freeze substitution was removed before warming began. The best ultrastructure was achieved by freeze substituting in acetone containing uranyl acetate at -90°, followed by gradual warming to 0° in pure acetone and rehydration between 0° and room temperature (Figure 1a). Rehydrated samples were processed for transmission EM at room temperature using our standard protocol (Ostroff et al., 2010), which includes two osmium staining steps and *en bloc* staining with uranyl acetate. This protocol consistently produced excellent ultrastructural preservation (Figure 1b), with intact membranes and fine ultrastructural features (Figure 1c). Inferior results were obtained when freeze substitution was performed in acetone alone or with glutaraldehyde or tannic acid instead of uranyl acetate.

**Figure 1.**
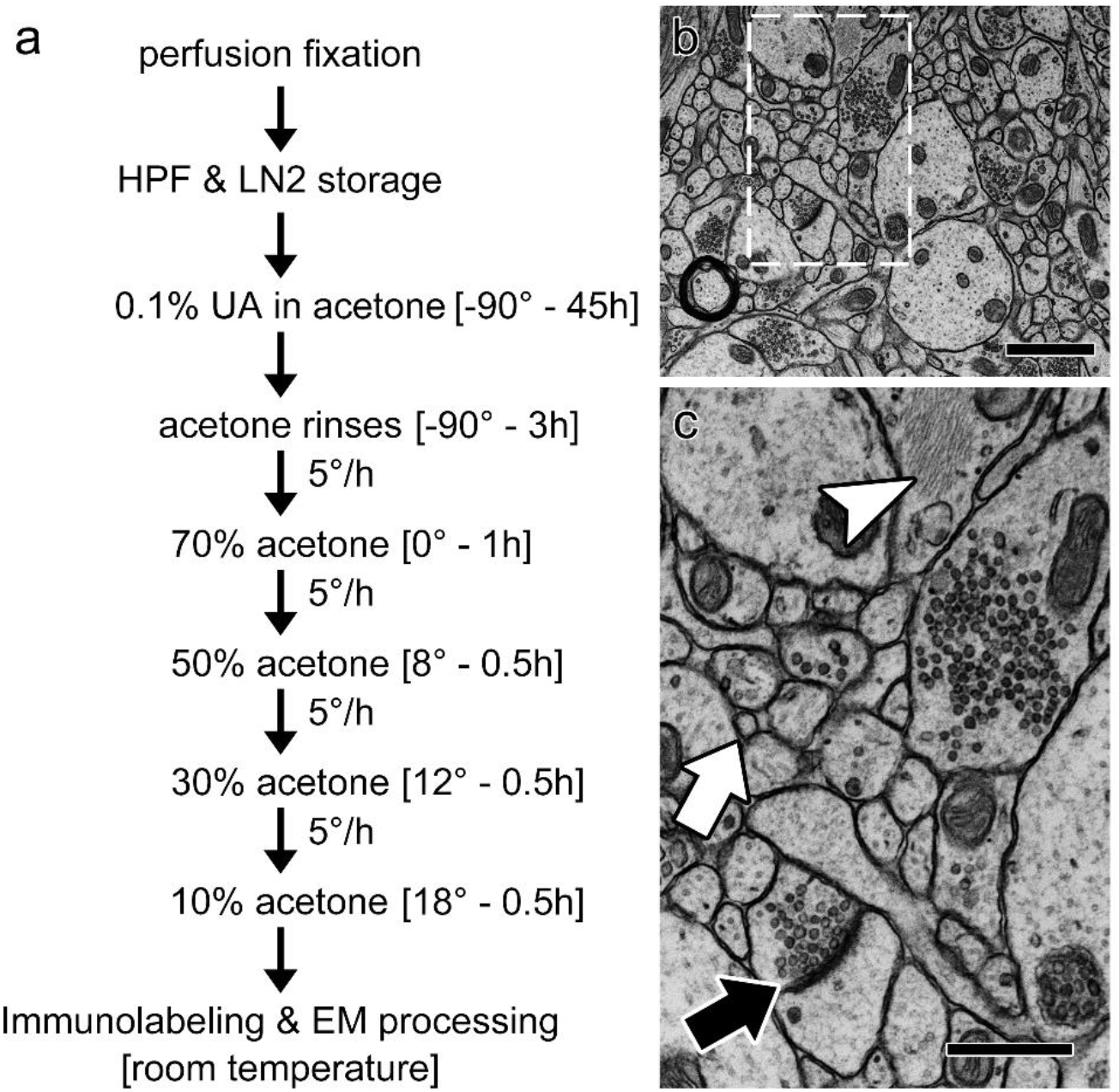
Rehydration protocol and ultrastructure. a) Rehydration schedule. HPF: high-pressure freezing; LN2: liquid nitrogen; UA: uranyl acetate. b) Electron micrograph of a rehydrated sample. c) Enlargement of the outlined region in (a) showing preservation of fine structures such as a spine neck (white arrow), a synapse (black arrow), and glial filaments (arrowhead). Scale bar = 1 µm in (a), 500 nm in (b).

### Preservation of ultrastructure and antigenicity in rehydrated tissue

Primary fixation with glutaraldehyde is considered optimal for preservation of brain tissue ultrastructure (Karnovsky, 1965; Schultz and Karlsson, 1965). Although glutaraldehyde is considered detrimental to immunolabeling due to extensive crosslinking and denaturation of some antigens, treatment with the reducing agent sodium borohydride and signal amplification with avidin-biotin complex (ABC) greatly enhance signal in glutaraldehyde-fixed neuronal tissue (Eldred et al., 1983; Willingham, 1983; Mrini et al., 1995). To verify that primary fixation in glutaraldehyde would permit detection of soluble cell-type markers, we compared labeling after perfusion with 4% paraformaldehyde alone or with the addition of 2.5% glutaraldehyde. Vibratome sections were treated with NaBH_4_ and antibody labeling was developed with ABC and the peroxidase substrate 3,3’- diaminobenzidine (DAB). Light microscopy revealed that labeling of neurons and processes was similarly dense in both fixatives for the calcium-binding proteins calbindin and parvalbumin, while the amino acid neurotransmitter GABA, as expected, required glutaraldehyde (Figure 2).

**Figure 2.**
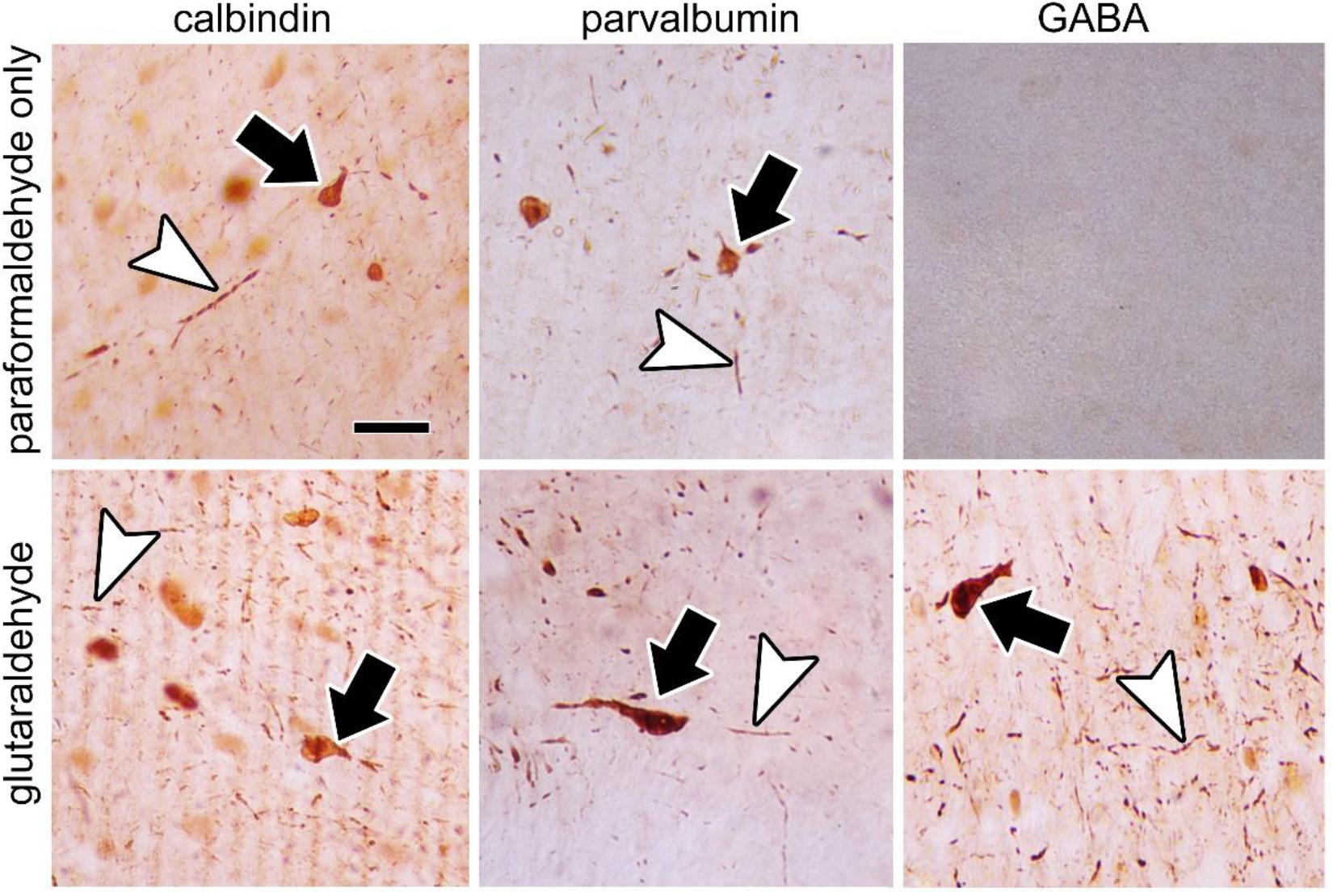
Immunolabeling in different primary fixatives. Rats were perfused with 4% paraformaldehyde without (top row) or with (bottom row) 2.5% glutaraldehyde before labeling with antibodies to calbindin, parvalbumin, or GABA. Examples of labeled cell bodies (arrows) and neuronal processes (arrowheads) are shown. Scale bar = 20 µm.

We next tested the performance of our rehydration protocol in pre-embedding immunolabeling and ultrastructural preservation. Vibratome sections from the same adult rat brains were either immunolabeled or high-pressure frozen on the day of sectioning. The frozen samples were stored in liquid nitrogen before freeze substitution, rehydration, and immunolabeling, and all samples were embedded for EM. Ultrathin sections from non-immunolabeled samples were stained with uranyl acetate and lead to enhance contrast (Figure 1b-c), but because DAB labeling can be obscured by on-section staining it was omitted for all further samples. Ultrastructural preservation in the labeled rehydrated samples was excellent, whereas broken membranes were found in the non-frozen samples (Figure 3).

**Figure 3.**
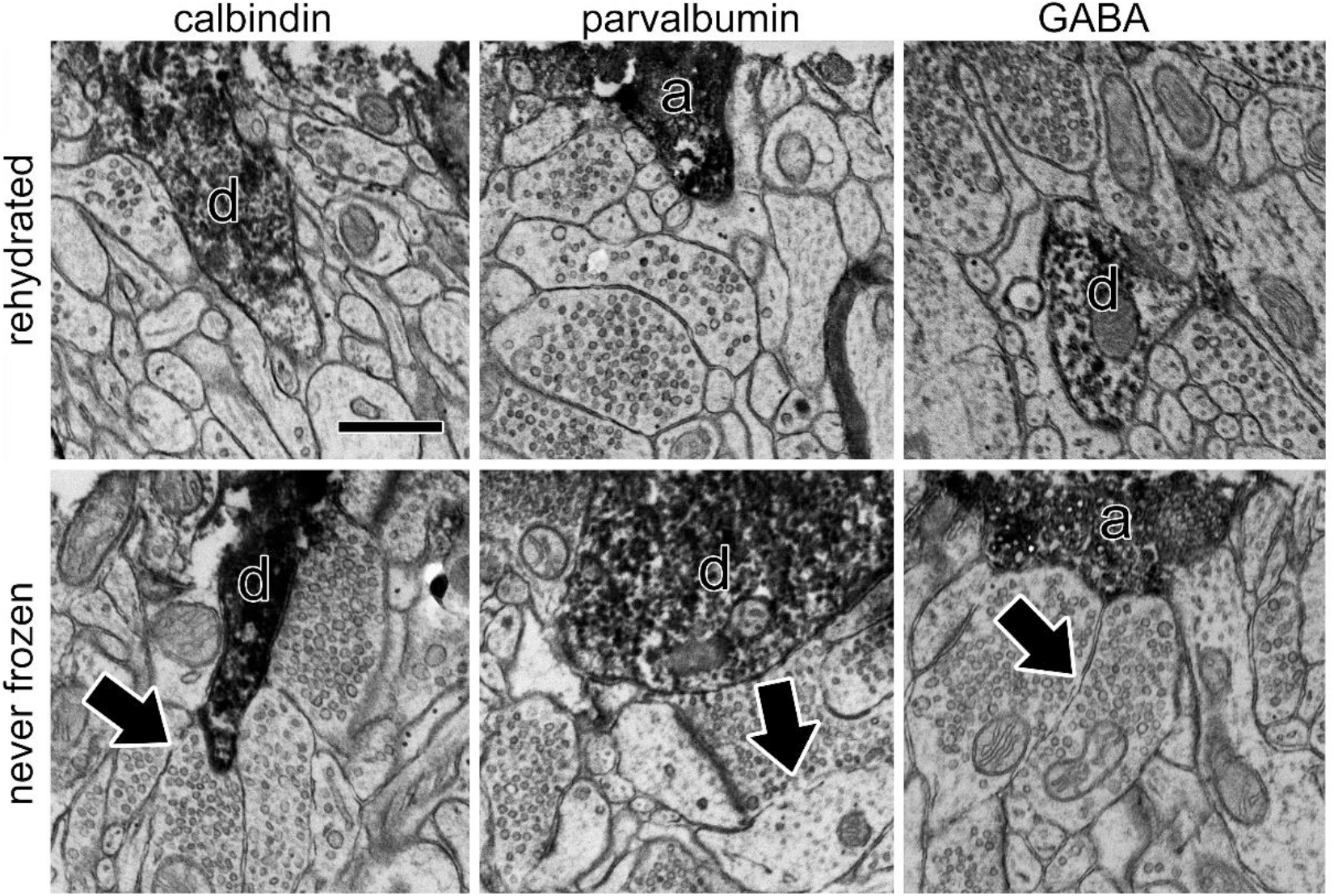
Immunolabeling of rehydrated samples. Tissue samples from the same rats were either high-pressure frozen and rehydrated before immunolabeling (top row) or labeled immediately after perfusion without freezing (bottom row). A labeled dendrite (d) or axon (a) is visible in each image. Broken membranes (arrows) are noticeable only in the samples that were not frozen and rehydrated. Scale bar = 500nm.

Labeled axons and dendrites were easily identifiable in both preparations.

### Detergent permeabilization of rehydrated tissue

The non-ionic detergent Triton-X 100 is widely used to permeabilize aldehyde-fixed tissue for immunohistochemistry at the light microscopy level, but its damaging effects on membrane ultrastructure limit its use in EM (Humbel et al., 1998). Although treatment with Triton-X 100 can redistribute proteins within fixed cells (Melan and Sluder, 1992) and reduce or alter labeling patterns for some antigens (McDonald and Mascagni, 2021), many antigens do label more strongly with Triton-X 100 (Sesack et al., 2006), sometimes even in a cell-type specific manner (Aoki et al., 1987). Because our rehydration protocol resulted in improved ultrastructure relative to standard EM processing, we wondered whether it would mitigate the effects of detergent treatment. Samples were prepared as above and subjected to one of three treatments: addition of 0.2% Triton-X 100 to the blocking buffer during immunolabeling, addition of 0.1% Triton-X 100 to the first step of the rehydration procedure (70% acetone at 0°), or no detergent. Treatment with Triton-X 100 at room temperature was highly destructive to membranes, but the lower concentration used at 0° resulted in EM morphology that was only slightly worse that that of non-permeabilized tissue (Figure 4).

**Figure 4.**
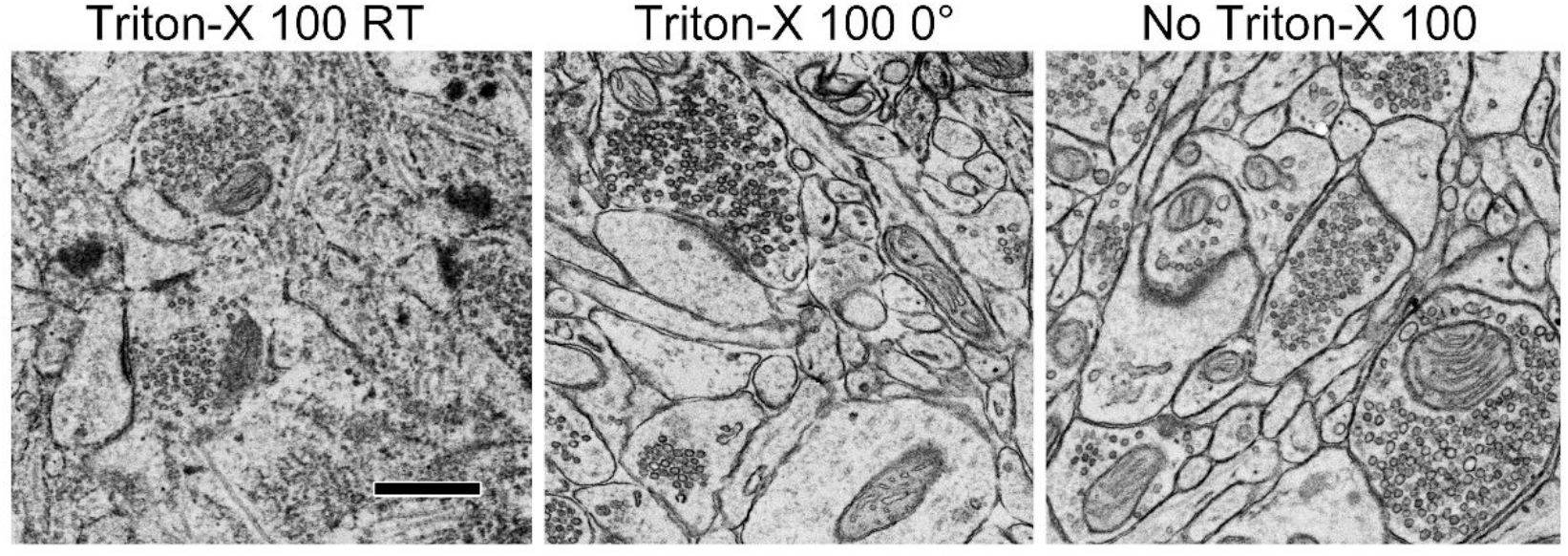
Effect of Triton-X 100 on rehydrated samples. EM images of rehydrated samples immunolabeled for either calbindin or GABA with Triton-X 100 in the room temperature blocking step at 0.2% (left), in the 0° rehydration step at 0.1% (center), or without Triton-X 100 (right). Scale bar = 500 nm.

### Ultrastructure is preserved throughout the depth of rehydrated tissue

EM volume reconstruction requires uniformly excellent morphology throughout a tissue sample so that neuronal processes can be reconstructed across long distances. To ensure that our rehydration protocol did not produce uneven results within sample blocks, we embedded tissue in flat molds and sectioned perpendicular to the vibratome edges (Figure 5a) so that the entire 100 µm thickness was visible on each section (Figure 5b). Ultrastructural preservation was of similarly high quality at the vibratome edges and in the center of the samples (Figure 5c). DAB labeling generally only penetrated 5 or 10 µm from the cut edges, which was unsurprising in well-preserved, non-permeabilized tissue. In the context of 3D reconstruction this is an advantage; chromogenic labels are sensitive but occlude subcellular structures, so limiting the label to the sample surface allows processes to be identified while leaving their structure largely free from debris.

**Figure 5.**
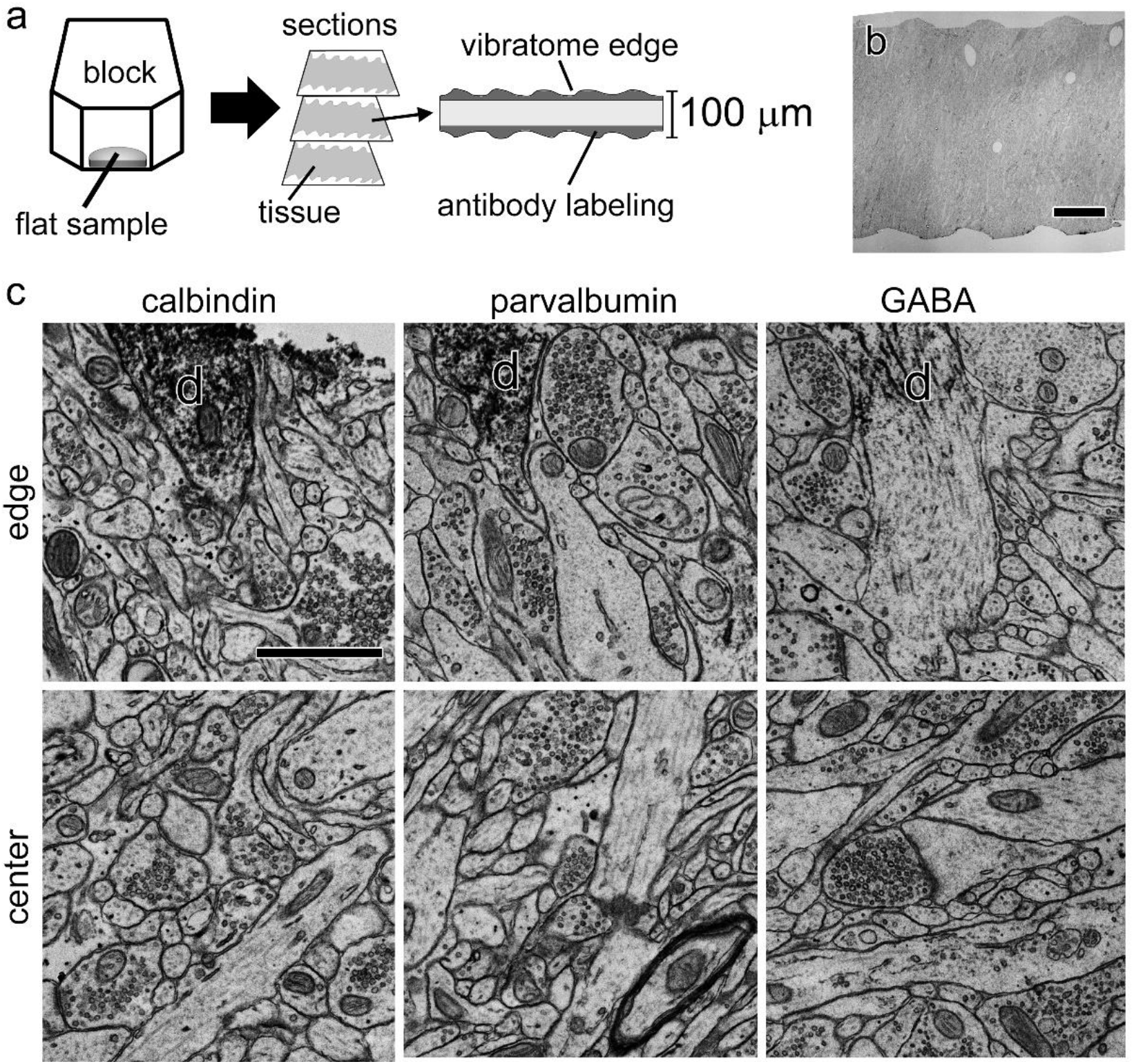
Labeling and morphology through sample thickness. a) Samples were embedded flat in resin blocks so that the place of section was perpendicular to the vibratome edges. Immunolabeling reagents penetrate only a few microns from each edge. b) Low-magnification EM image showing the full 100µm sample thickness. c) Example EM images of immunolabeled samples taken close to the vibratome edge (top row) or in the center of the sample depth (bottom row). Labeled dendrites (d) are visible only at the edges. Scale bar = 25µm in (b), 1µm in (c).

### Rehydration results are reproducible

Ice crystal damage or solvent extraction could easily render samples unusable for serial reconstruction, and it is therefore essential that a rehydration protocol produce reliably good ultrastructure. Only a handful of studies have reported rehydration after freeze substitution (Robinson and Karnovsky, 1991; van Donselaar et al., 2007; Ripper et al., 2008; Hess et al., 2018; Tsang et al., 2018) and just one involved brain tissue (Tsang et al., 2018), so the reproducibility of rehydration is unknown. To ensure that rehydration could be safely relied upon for use on irreplaceable samples, we performed the procedure over a dozen times using samples from several different rats. We saw no evidence of ice damage in any sample. A few samples were judged to have lower morphological quality, but these were from a single rat and other samples in the same rehydration experiments were well-preserved, meaning that the cause was poor primary fixation (likely sub-optimal perfusion) and not rehydration. Figure 6 shows representative images from four different blocks taken from three different rats and three rehydration experiments.

**Figure 6.**
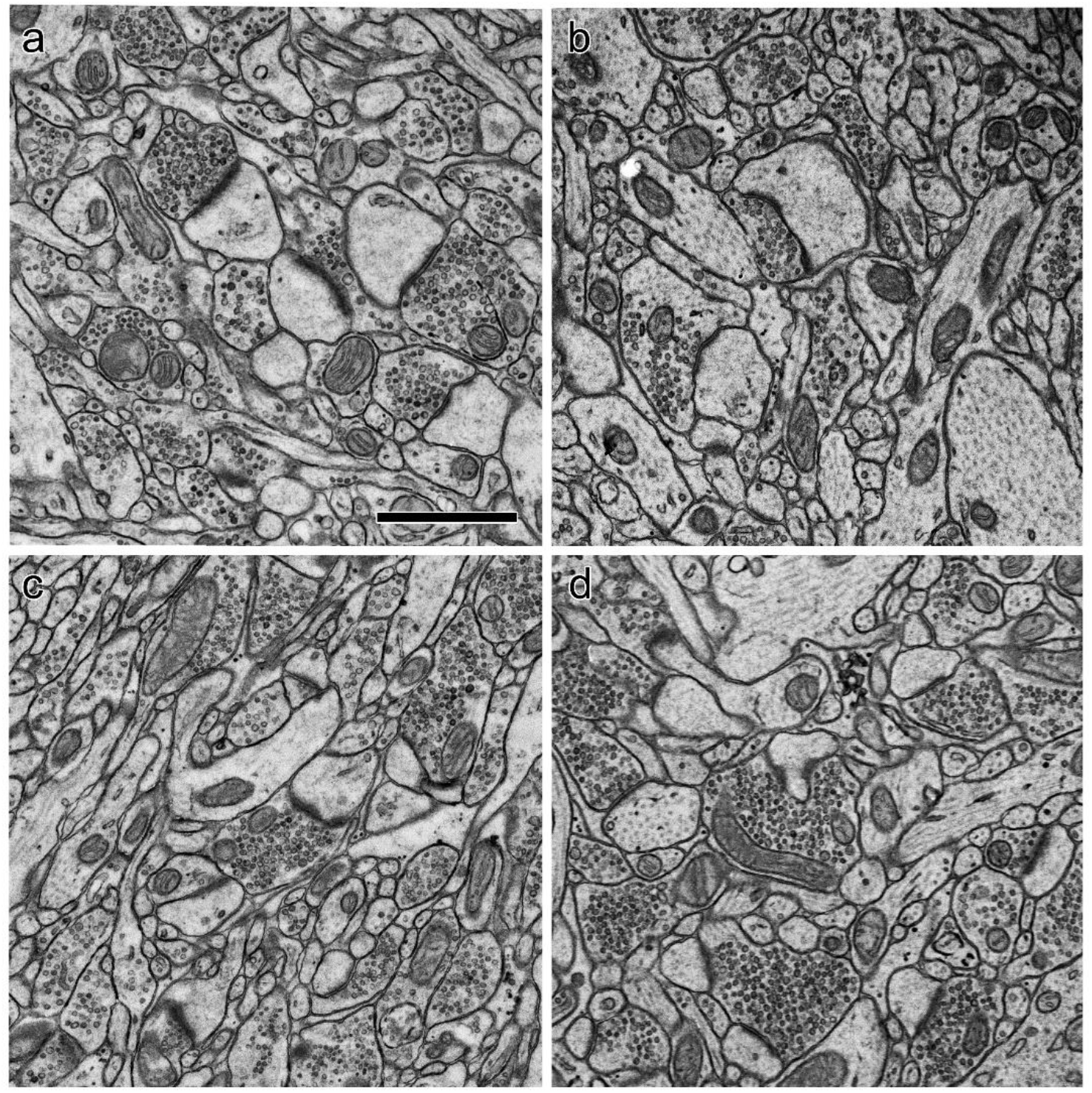
Reproducibility of ultrastructure in rehydrated samples. a-b) EM images of rehydrated samples from two different rats collected and rehydrated in different experiments. c-d) Two separate samples from a third rat from a different rehydration experiment. Scale bar = 1 µm.

## Discussion

Pre-embedding immunolabeling is much more sensitive and reliable than post-embedding for most antigens (Polishchuk and Polishchuk, 2019), and neuroanatomists have a long history of using pre-embedding labeling to study the ultrastructure of molecularly-defined synapses (Sesack et al., 2006). Serial section EM reconstruction provides much richer data than single-section analysis of brain tissue, and pre-embedding labeling of endogenous molecules can greatly enhance the value of these datasets by adding cell-type identification (Zikopoulos et al., 2016) and revealing the subcellular distributions of molecules (Gindina et al., 2021). The compromised ultrastructural preservation inherent in pre-embedding labeling, however, is a deterrent to routine use in serial reconstructions.

We have developed a reliable, reproducible protocol for preventing morphological damage during immunolabeling, eliminating the trade-off between efficient serial section reconstruction and identification of endogenous molecules.

EM reconstruction of brain tissue requires fully intact, highly contrasted membranes so that small neuronal processes can be followed across large numbers of serial sections. Standard pre-embedding antibody labeling procedures can damage membrane integrity in multiple ways. Membranes can be intentionally damaged with detergents, freeze-thaw cycles, or bacterial toxins to create holes for antibodies to pass through (Eldred et al., 1983; Pickel et al., 1986; Humbel et al., 1998).

Aside from actively damaging membranes, failure to preserve them can also degrade tissue quality. High concentrations of glutaraldehyde in the primary fixative benefit membrane integrity (Karnovsky, 1965; Schultz and Karlsson, 1965), but the resulting dense crosslinks impede antibody penetration. Lower concentrations or alternatives such as acrolein are commonly used to enhance immunolabeling, but at some cost to ultrastructure (Sesack et al., 2006). Extensive antibody incubation times are also used to enhance penetration (Fulton and Briggman, 2021), but delaying lipid fixation with osmium tetroxide while antibody incubations are performed can degrade membrane structure. Our rehydration protocol includes freeze substitution in uranyl acetate, which acts as a lipid fixative and preserves antigenicity in cryoembedded brain tissue (Erickson et al., 1987; van Lookeren Campagne et al., 1991; Giddings, 2003). It is likely that membrane fixation by uranyl acetate is responsible for the improved ultrastructure we observe in rehydrated samples relative to samples immunolabeled immediately after perfusion. Uranyl acetate can provide significant membrane stabilization in the absence of osmium (Berryman and Rodewald, 1990; Phend et al., 1995; Burette et al., 2012), and was used in a previous report of osmium-free rehydration (Tsang et al., 2018).

Because a delay between aldehyde fixation and osmium fixation can degrade ultrastructure, pre-embedding labeling must be performed at the time that samples are collected. This is not always convenient, for example if large numbers of samples must be collected at once or if sample availability is unpredictable. It also means that antibodies and labeling protocols must be chosen and validated at the time of sample collection, which limits future options for a given sample. Unlike post-embedding labeling, where many antibodies can be applied to separate ultrathin sections (Anderson et al., 2011; Shahidi et al., 2015), pre-embedding labeling is generally limited to combinations of no more than three peroxidase substrates or gold labels (Sesack et al., 2006; Polishchuk and Polishchuk, 2019). For valuable samples, it could be advantageous to dissect smaller pieces for later use with different antibodies. A major advantage of HPF is that it allows samples to be stored indefinitely under liquid nitrogen, so by making HPF compatible with pre-embedding immunolabeling our protocol enables much greater flexibility in experimental design.

Despite its higher sensitivity, pre-embedding is considered inferior to post-embedding in many EM applications because of limited antibody penetration. Labeling depth can be greatly enhanced by using genetically-encoded enzymes which do not require bulky reagents for visualization (Schikorski et al., 2007; Martell et al., 2012; Cruz-Lopez et al., 2018). These enzymes have been used in EM reconstructions of *Drosophila* sensory neurons after HPF and rehydration (Zhang et al., 2019; Gonzales et al., 2021), but their applications are limited because they require transgenesis and do not detect endogenous molecules. Although limited label penetration is a problem for quantitative molecular localization studies and sparse antigens, in the context of serial section reconstruction of brain circuits it can be considered an advantage. Peroxidase substrates obscure ultrastructural details, as do gold labels to a lesser extent, so restricting labeling to a few microns near the cut surface of a process enables molecular identification at the edge as well as careful morphological analysis of the unlabeled interior (Zikopoulos et al., 2016). By allowing high-pressure freezing to be used with sensitive pre-embedding labeling and aqueous heavy metal staining, our protocol should eliminate the major barrier to identifying endogenous cell type markers in EM reconstructions of brain tissue.

## Materials & Methods

### Subjects

Subjects were adult (8-12 weeks) female Sprague-Dawley rats (Hilltop, Scottdale, PA). Rats were pair housed on a 12-hour reverse light/dark cycle with *ad libitum* food and water. All animal procedures were approved by the Animal Care and Use Committee of the University of Connecticut.

### Perfusion and high-pressure freezing

Rats were deeply anesthetized with chloral hydrate (750mg/kg) and transcardially perfused with 500 ml of fixative using a peristaltic pump at a rate of 75 ml/min. The fixative consisted of 2.5% glutaraldehyde and 4% freshly depolymerized paraformaldehyde with 4 mM MgCl_2_ and 2 mM CaCl_2_ in 0.1 M PIPES buffer at pH 7.4. Aldehydes and buffer were obtained from Electron Microscopy Sciences (Hatfield, PA) and salts from Sigma-Aldrich (St. Louis, MO). Brains were removed immediately and immersed in the perfusion fixative for one hour, then rinsed in the perfusion buffer and sectioned at 100 µm on a vibrating slicer (Leica Biosystems). All steps were carried out at room temperature. The area around the LA was dissected using 2 mm biopsy punch.

For high-pressure freezing, samples were loaded into aluminum carriers with 20% polyvinylpyrrolidone (EMD Millipore Corp., Burlington, MA) as a filler, then frozen in a Wohlwend Compact 3 high-pressure freezer (Technotrade International, Inc., Manchester, NH). Samples were stored under liquid nitrogen until further processing.

### Freeze substitution and rehydration

For freeze substitution, cryotubes were filled with 0.1% uranyl acetate (SPI supplies., West Chester, PA) in acetone and frozen by immersion in liquid nitrogen.

Samples were placed atop the frozen solution and the cryotubes were transferred to an AFS2 freeze substitution unit (Leica Microsystems). Samples were held at -90° for 45 hours, then the substitution medium was replaced with three changes of pure acetone over three hours. The temperature was then raised to 0° at a rate of 5° per hour before rehydration began. The acetone was then replaced with an ascending sequence of acetone dilutions in water. Samples were incubated for one hour in 70% acetone at 0°, then for 30 minutes in 50%, 30%, and 10% acetone at 8°, 12°, and 18° respectively. Warming between steps was performed at 5° per hour. At the end of the 10% acetone step samples were rinsed in 0.1 M PIPES buffer at room temperature. For tests of low-temperature permeabilization, 0.1% Triton-X 100 (Sigma-Aldrich) was added to the 70° acetone solution.

### Immunolabeling

All immunolabeling was performed at room temperature in 0.1 M PIPES buffer, pH 7.4. Samples were first reacted with 1% sodium borohydride (Sigma-Aldrich) for 15 minutes, rinsed in buffer, and incubated in 0.3% hydrogen peroxide for 15 minutes to quench endogenous peroxidases. They were then blocked for one hour in 1% bovine serum albumin (Jackson Immuno Research Laboratories, West Grove, PA). Triton-X 100 was added at 0.2% during this step for tests of permeabilization at room temperature. Samples were incubated overnight in primary antibody overnight, then rinsed and incubated for one hour in secondary antibody. Labeling was detected using an avidin/biotin peroxidase kit (Vectastain Elite ABC-HRP kit PK-6100, Vector Laboratories, Burlingame, CA) according to the manufacturer’s instructions, followed by reaction with 1 mM 3,3’- diaminobenzidine (Sigma-Aldrich) in 0.003% H_2_O_2_ for 8 minutes.

### Antibodies

The following primary antibodies and dilutions were used: Sigma-Aldrich a2052 rabbit anti-GABA at 1:10,000; Synaptic Systems 214 003 rabbit anti-calbindin at 1:500; Abcam ab11427 rabbit anti-parvalbumin at 1:500. The secondary antibody for all experiments was Invitrogen 65-6140 goat anti-rabbit biotin conjugate at 1:200.

### Electron and light microscopy

Processing for EM was as previously described (Ostroff et al., 2010). Briefly, samples were rinsed in 0.1 M cacodylate buffer, pH 7.4, post-fixed in 1% osmium tetroxide with 1.5% potassium ferrocyanide followed by 1% osmium tetroxide, dehydrated in an ascending series of ethanol dilutions containing 1.5% uranyl acetate, and flat embedded in LX-112 epoxy resin (Ladd Research Industries, Williston, VT). Blocks were trimmed to expose a region at the center of the dorsolateral LA, and sections were cut 45 nm on a Leica UC7 ultramicrotome and collected on pioloform-coated slot grids. Imaging was performed at 6000X on a JEOL 1400 transmission EM with an AMT Nanosprint-43 Mark II digital camera (Advanced Microscopy Techniques, Woburn, MA). The sections shown in Figure 1a-b were stained with saturated aqueous uranyl acetate and Sato’s lead (Hanaichi et al., 1986); all other images are of unstained sections. For light microscopy, immunolabeled samples were mounted in DPX and imaged at 40X on an upright microscope using an SLR camera (Canon). Images were cropped and contrast adjusted using Photoshop software (Adobe).

## Acknowledgements

We are grateful to Maritza Abril for expert technical assistance and to Mark Terasaki for helpful comments on the manuscript. This work was performed in part at the Bioscience Electron Microscopy Laboratory of the University of Connecticut.

